# Three R2R3-MYB transcription factors from banana (*Musa* spp.) activate structural anthocyanin biosynthesis genes as part of an MBW complex

**DOI:** 10.1101/2022.08.15.503939

**Authors:** Mareike Busche, Boas Pucker, Bernd Weisshaar, Ralf Stracke

## Abstract

Bananas are among the most popular fruits in the world and provide food security and employment opportunities in several developing countries. An increased anthocyanin content could enhance the health promoting properties of banana fruits. The biosynthesis of anthocyanins is largely regulated at the transcriptional level. However, little is known about transcriptional activation of anthocyanin biosynthesis in banana. We analysed the regulatory activity of three *Musa*MYBs predicted by bioinformatic analysis to transcriptionally regulate anthocyanin biosynthesis in banana. *MusaMYBA1, MusaMYBA2* and *MusaMYBPA2* did not complement the anthocyanin deficiency phenotype of the *A. thaliana pap1/pap2* mutant. However, co-transfection experiments in *A. thaliana* protoplasts showed that *Musa*MYBA1, *Musa*MYBA2 and *Musa*MYBPA2 function as components of a transcription factor complex with a bHLH and WD40 protein, called MBW complex, resulting in the activation of the *anthocyanin synthase* and *dihydroflavonol 4-reductase* promoters from *A. thaliana*. The activation potential of *Musa*MYBA1, *Musa*MYBA2 and *Musa*MYBPA2 increased when combined with the monocot bHLH *Zm*R instead of the dicot *At*EGL3. This work paves the path towards decoding the MBW complex-mediated transcriptional activation of anthocyanin biosynthesis in banana. Moreover, it facilitates research towards an elevated anthocyanin content in banana and other monocot crops.

## Introduction

Bananas (*Musa*) are monocotyledonous, perennial plants which are grown in many tropical and subtropical countries. They are one of the most important food crops, particularly in the developing world (1). While the sweet fruits of dessert bananas are popular in Europe and Northern America, plantains or cooking bananas are commonly eaten as a staple food in Africa and Latin America where they provide food security, as well as employment opportunities (2). Furthermore, banana fruits are rich in several health promoting minerals and beneficial phytochemicals such as vitamins and flavonoids (3).

Flavonoids are a major group of plant specialised metabolites which share a basic structure of two aromatic C6-rings connected by a heterocyclic ring (4). Reorganisations and modifications e.g. oxidation, glycosylation, acylation, and methylation of the carbon skeleton create a versatile group comprising more than 9,000 different flavonoid derivatives (5). Consequently, flavonoids do not only contribute to the nutritional value of fruits, but also play important roles in manifold processes. While the group of red coloured anthocyanin pigments attracts animals for pollination and dispersal of seeds by colouring flowers and fruits, other flavonoids protect plants against UV-B irradiation or increase plant fertility (6-10). Flavonoids from many species have been reported to have anti-pathogenic properties, this includes flavonoids from *Dianthus caryophyllus* which exhibit antifungal activity against the plant’s major pest *Fusarium oxysporum* f.sp. *dianthi* (11, 12). The tropical race 4 (TR4) of the banana Fusarium wilt (commonly known as ‘panama disease’) is caused by another *Fusarium* subspecies called *Fusarium oxysporum* f. sp. *cubense* (Foc) and threatens the global banana production (13). Transcriptome analysis of susceptible and resistant banana cultivars infected by Foc TR4 revealed an increased transcription of flavonoid biosynthesis related genes in the resistant cultivar, suggesting an involvement of flavonoids in the defence against Foc TR4 (14).

Flavonoid biosynthesis is one of the best characterised pathways of the secondary metabolism and has been extensively studied in many plant species (15). In banana, several flavonoid biosynthesis related enzymes have been identified and characterised (16, 17). Regulation of structural genes on a transcriptional level allows a specific response to environmental influences as well as development and organ specific expression (18-20). MYB transcription factors are common transcriptional regulators of flavonoid biosynthesis. While some MYBs act independently, others interact with basic helix-loop-helix (bHLH) and WD40 proteins to form a protein complex called MBW complex (21). MYB proteins are present in all eukaryotes and characterised by highly conserved DNA-binding domains (22). These MYB domains consist of up to three imperfect amino acid repeat sequences, based on which they are classified. R2R3-MYBs are the most abundant class of plant MYBs and reveal versatile functions in plant-specific processes (23). Besides core- and specialised metabolism they are also involved in cell fate and -identity definition, developmental processes and the response to biotic and abiotic stresses (23). Well-known R2R3-MYBs which act as activators of anthocyanin biosynthesis include C1 (COLOURED ALEURONE1) from *Zea mays* (*Z. mays*), as well as PAP1 (PRODUCTION OF ANTHOCYANIN PIGMENT1/MYB75) and PAP2 (MYB90) from *Arabidopsis thaliana* (*A. thaliana*) (24, 25). They act as part of an MBW complex and control the promoters of the anthocyanin biosynthesis related structural genes as for example *ANTHOCYANIDIN SYNTHASE* (*ANS*) and *DIHYDROFLAVONOL 4-REDUCTASE* (*DFR*) (26-29).

In banana (*Musa acuminata, M. acuminata*), 285 R2R3-MYB proteins, including several putative regulators of flavonoid biosynthesis, have been identified in a genome-wide study (30). Furthermore, MYB31, MYB4 and MYBPR1 - MYBPR4 have been identified as negative regulators of flavonoid biosynthesis in banana (31, 32). Despite the recent identification of two proanthocyanidin biosynthesis activating R2R3-MYBs (33), little functional data is available on positive regulators (activators) of flavonoid and in particular anthocyanin biosynthesis in *M. acuminata*.

Here, we describe the regulatory properties of three *Musa*MYBs, named *Musa*MYBA1, *Musa*MYBA2 and *Musa*MYBPA2, with a possible role in the regulation of anthocyanin biosynthesis. As one of these *Musa*MYBs was very recently published under the term *Musa*MYBPA2 (33), we used this name to avoid confusion due to multiple protein naming. Regulatory activity was investigated by *in planta* complementation experiments of the anthocyanin deficient *A. thaliana* regulatory *pap1/pap2* mutant and co-transfection experiments in *A. thaliana* protoplasts (for detailed methods refer to Supplementary File 2). Our results show that *Musa*MYBA1, *Musa*MYBA2 and *Musa*MYBPA2 are able to activate the promoters of *AtANS* and *AtDFR* as part of an MBW complex. Moreover, we show that the activation potential of *Musa*MYBA1, *Musa*MYBA2 and *Musa*MYBPA2 is increased when combined with the monocot bHLH *Zm*R instead of the dicot bHLH protein ENHANCER OF GLABRA 3 (*At*EGL3).

## Main text

We aimed to analyse the regulatory properties of three *Musa*MYBs which have been previously assigned with a possible role in positive regulation of anthocyanin biosynthesis (Ma06_g05960 or *MusaMYBA1*, Ma09_g27990 or *MusaMYBA2*, Ma10_g17650 or *MusaMYBPA2*). Since all three *MusaMYB* genes were detected in the haploid *M. acuminata* reference genome sequence DH (doubled-haploid) Phang v2 (34, 35), these *MYBs* appear to be present in the same sub-genome, suggesting that they are different genes and not haplo-copies. We tried to amplify the corresponding coding sequences (CDSs) on a template collection containing cDNA from different banana samples. The CDSs of all three *MusaMYB*s were successfully amplified on cDNA derived from peel tissue.

In a first approach, we performed a complementation assay with the regulatory *A. thaliana pap1/pap2* double mutant which cannot produce anthocyanins in the seedling (Figure 1). Seedlings were grown on anthocyanin synthesis-inducing media to analyse the ability of 2×35S-driven *MusaMYB*s to complement the *pap1/pap2* anthocyanin deficiency. While wild-type seedlings accumulated high amounts of red anthocyanin pigments, *pap1/pap2* seedlings did not. Even though *MusaMYBA1, MusaMYBA2* or *MusaMYBPA2* were successfully expressed in the transgenic seedlings (Supplementary Figure S1), the anthocyanin level in *pap1/pap2* plants expressing *MusaMYBA1, MusaMYBA2* or *MusaMYBPA2* did not differ from the double mutant. Accordingly, *MusaMYBA1, MusaMYBA2* and *MusaMYBPA2* do not seem to be able to complement the mutant phenotype and thus to regulate anthocyanin biosynthesis in *A. thaliana* in combination with the *bHLH* and *WD40* genes expressed in *A. thaliana* seedlings.

**Figure 1:**
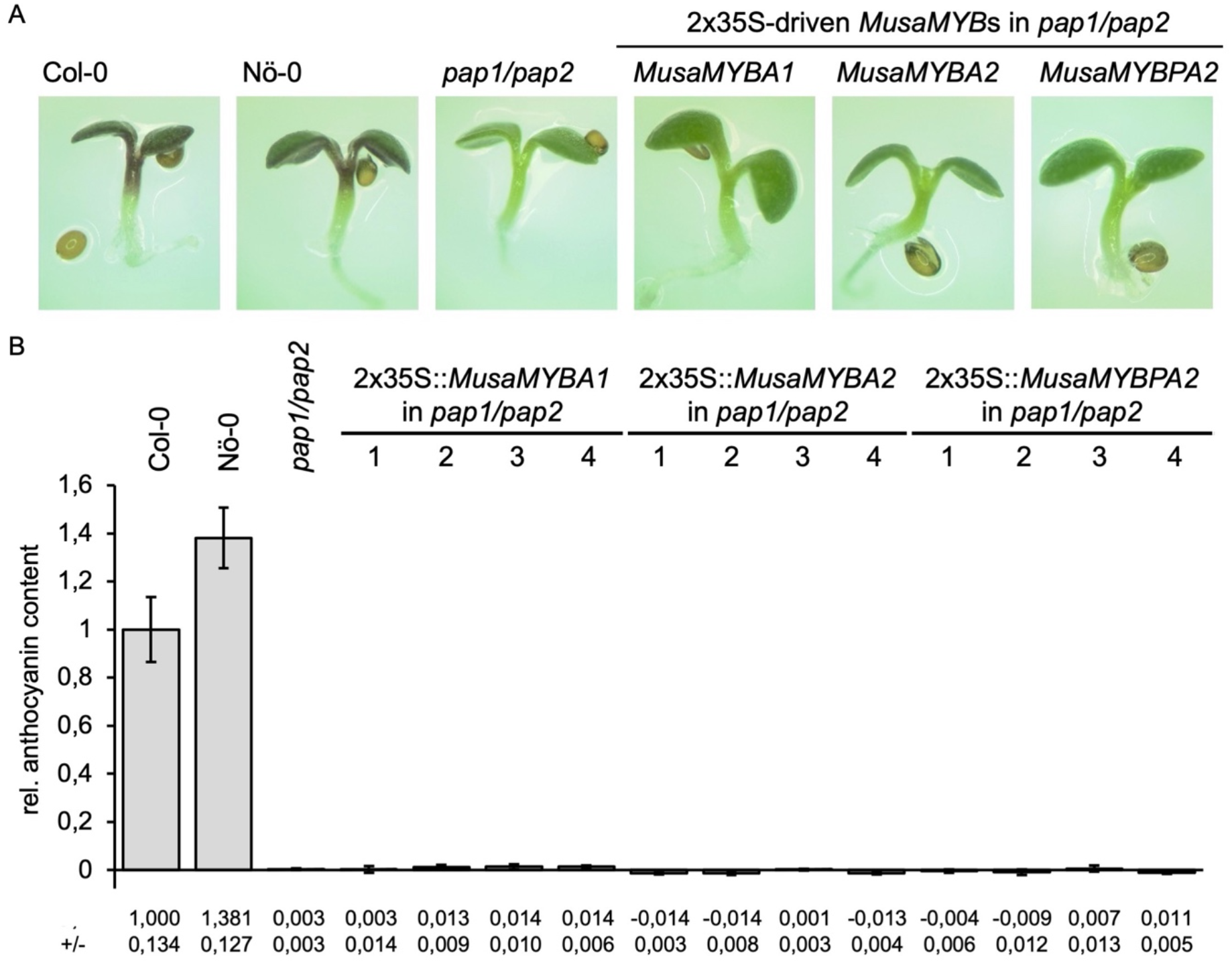
*MusaMYBA1, MusaMYBA2* and *MusaMYBPA2* cannot complement the anthocyanin deficient phenotype of *A. thaliana pap1/pap2* mutant seedlings. (A) Representative pictures of anthocyanin accumulation in 6-day-old *MusaMYB*-expressing *pap1/pap2* seedlings. A representative plant per construct is displayed. (B) Photometrical measurement of the sucrose induced anthocyanin content in *pap1/pap2* seedlings expressing 2×35S-driven *MusaMYB*s. Col-0, Nö-0 (wildtypes) and *pap1/pap2* were used as controls. Error bars indicate the standard deviation of three biological replicates. The different numbers represent individual, independent, transgenic lines

Lloyd *et al*. (28) analysed the *Z. mays* anthocyanin regulator C1 by generation of transgenic *A. thaliana* overexpression lines. In their study, Lloyd *et al*. generated three independent *ZmC1*-expressing *A. thaliana* lines that did not show an increased anthocyanin content compared to wildtype. Similar results were obtained in transgenic tobacco. However, further experiments suggested that *Zm*C1 needs to interact with the maize bHLH *Zm*R to activate anthocyanin biosynthesis in the heterologous *A. thaliana* system (28). Phylogenetic analysis (36) showed that the anthocyanin biosynthesis regulating R2R3-MYBs from several monocots such as *Z. mays* and *Oryza sativa* (*O. sativa*) are part of a different phylogenetic clade than the anthocyanin regulators from several dicots, such as *A. thaliana* or *Vitis vinifera* and further monocots, including *Allium cepa* and *Lilium hybrida*. Experiments in snapdragon also showed that the monocot *Ac*MYB1 can activate anthocyanin production in dicots (36). The phylogenetic differences between anthocyanin biosynthesis activating R2R3-MYBs and the dependency of *Zm*C1 on *Zm*R in dicots hint at an explanation for our observations. For example, the *Musa*MYBs could depend on their endogenous or at least monocot bHLH for effective activation of structural anthocyanin biosynthesis genes. To investigate the phylogenetic differences between anthocyanin biosynthesis regulating R2R3-MYBs from different plant species, an approximately maximum-likelihood tree was built (Figure 2A). The resulting tree, which also contained R2R3-MYBs which activate other branches of flavonoid biosynthesis, revealed two distinct clades of anthocyanin-related R2R3-MYBs (highlighted in red). *Musa*MYBA1, *Musa*MYBA2 and *Musa*MYBPA2 form a clade with MYB10 from *Triticum aestivum* and exhibit a close evolutionary relationship with *Zm*C1, *Os*C1 and anthocyanin regulating MYBs from further monocots. In contrast, *At*PAP1 and *At*PAP2 fall into a second clade of anthocyanin-related MYBs. Furthermore, differences between anthocyanin biosynthesis regulating bHLHs were analysed in a second approximately maximum-likelihood tree (Figure 2B). The tree showed that monocot bHLH proteins involved in the regulation of anthocyanin biosynthesis form a separate clade (highlighted in red). Both, *Zm*R and several putative anthocyanin biosynthesis regulating bHLHs from banana fall into this clade. These phylogenetic analyses show that anthocyanin biosynthesis regulating R2R3-MYB and bHLH proteins from several monocot species seem to be distinct from other R2R3-MYB and bHLH proteins regulating anthocyanin biosynthesis, for example from *A. thaliana*. These differences could imply a dependency of *Musa*MYBA1, *Musa*MYBA2 and *Musa*MYBPA2 on a banana or other monocot bHLH.

**Figure 2:**
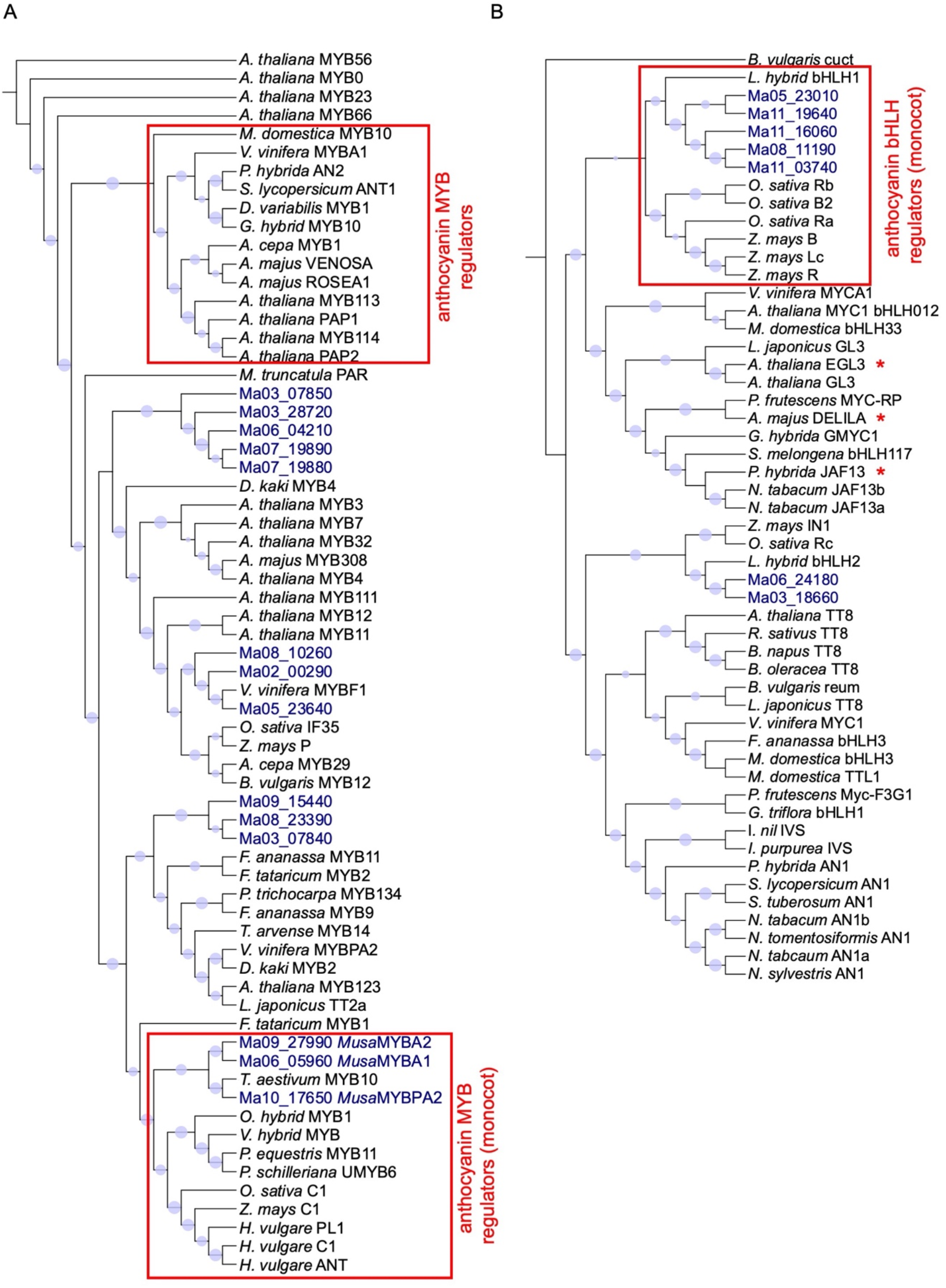
Rooted approximately maximum-likelihood trees of MYB (A) and bHLH (B) transcription factors. Circle sizes represent bootstrap values. Banana proteins are highlighted in blue.

To follow up this idea, we further investigated the regulatory properties of the three *Musa*MYBs performing co-transfection assays (Figure 3) in At7 protoplasts with different bHLH proteins from *A. thaliana* (*At*EGL3) or *Z. mays* (*Zm*R) and the *A. thaliana* WD40 protein TRANSPARENT TESTA GLABRA 1 (*At*TTG1). Thereby, their potential to activate the promoters of *AtDFR* and *AtANS*, which are important structural genes of anthocyanin biosynthesis (37), was analysed. While none of the *Musa*MYBs was able to independently activate *proAtANS* or *proAtDFR*, which both contain conserved *cis*-regulatory elements, a slight activation of *proAtDFR* was detectable when *Musa*MYBA1 or *Musa*MYBPA2 were combined with *At*EGL3 and *At*TTG1. In combination with *At*EGL3 and *At*TTG1, *Musa*MYBPA2 showed the strongest activation of *proAtDFR* and was also able to activate *proAtANS*. When combined with *Zm*R and *At*TTG1, the three MYBs *Musa*MYBA1, *Musa*MYBA2, and *Musa*MYBPA2 exhibited a significant activation potential on *proAtDFR* and *proAtANS*. All three *Musa*MYBs showed a higher activation potential in an MBW complex with *Zm*R and *At*TTG1 than in combination with *At*EGL3 and *At*TTG1.

**Figure 3:**
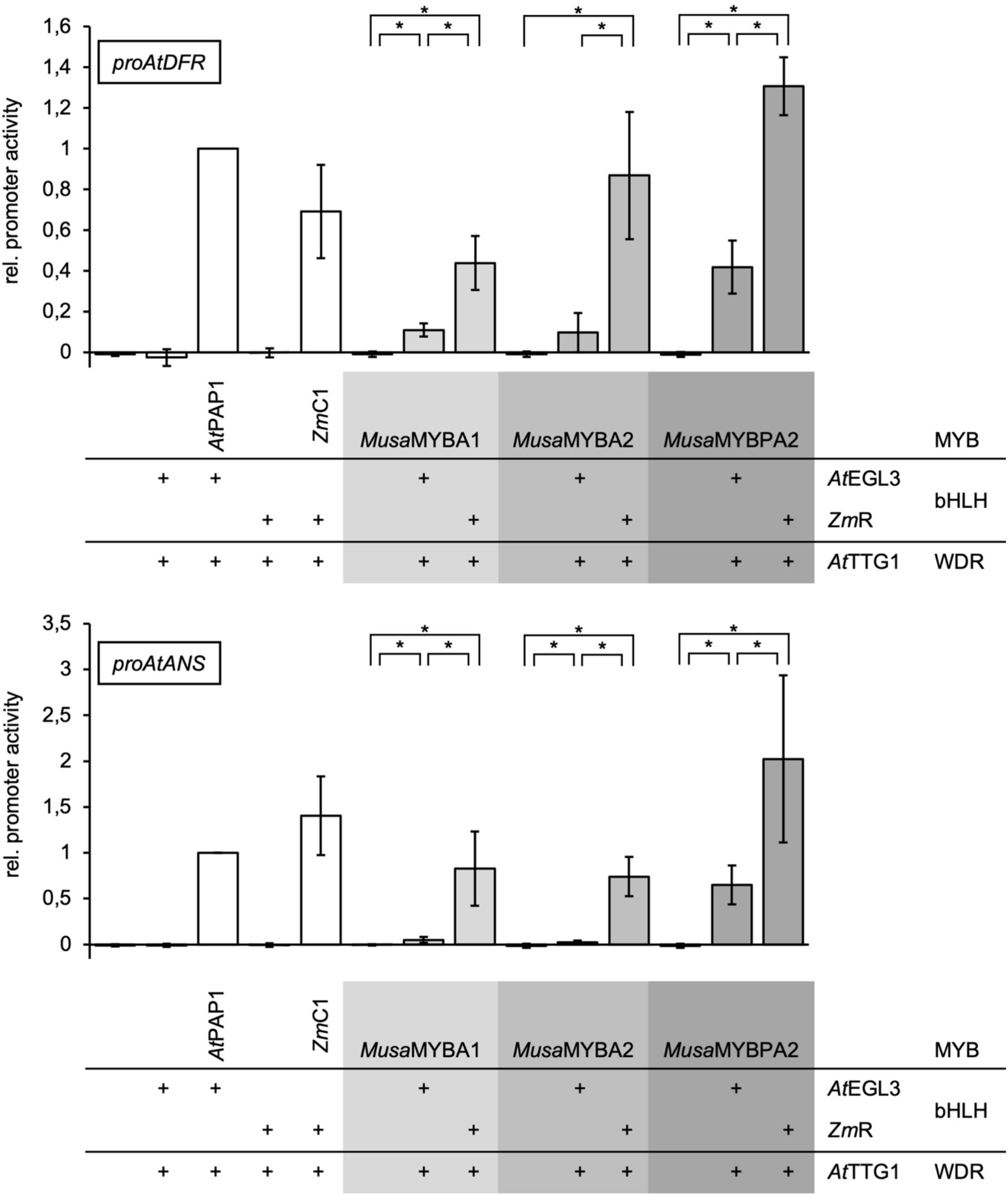
*Musa*MYBA1, *Musa*MYBA2 and *Musa*MYBPA2 can activate *proAtDFR* and *proAtANS* as part of an MBW complex. The ability of *Musa*MYBs to activate *proAtDFR-GUS* and *proAtANS-GUS* reporter constructs in combination with different bHLH proteins (*At*EGL3, *Zm*R) and a WDR (*At*TTG1) was analysed by co-transfection in *A. thaliana* At7 protoplasts. The relative promoter activity refers to the measured GUS reporter enzyme activity. Promoter activity is given relative to the values obtained for the *A. thaliana* MBW complex (*At*PAP1, *At*EGL3, *At*TTG1). Error bars indicate the standard deviation of five independent biological replicates. Statistical significance is indicated by asterisks which mark p-values < 0.05.

These results display that *Musa*MYBA1, *Musa*MYBA2 and *Musa*MYBPA2 are able to activate *proAtANS* and *proAtDFR* as part of an MBW complex and *Musa*MYBPA2 shows the strongest activation potential. Furthermore, *Musa*MYBA1, *Musa*MYBA2 and *Musa*MYBPA2 show a higher activation potential when combined with the monocot bHLH *Zm*R instead of the dicot bHLH *At*EGL3.

The previously mentioned phylogenetic differences could be an explanation for the observed higher activation potentials of the analysed *Musa*MYBs in combination with *Zm*R and *At*TTG1 instead of *At*EGL3 and *At*TTG1. It is conceivable that these differences impede the interaction between the tested *Musa*MYBs and the dicot bHLH *At*EGL3. As *Musa*MYBPA2 shows the highest activation potential and is also able to activate *proAtANS* in combination with *At*EGL3 or *Zm*R, differences between the three *Musa*MYBs that affect the interaction with the *At*EGL3 are likely and should be further studied. Interestingly, *Musa*MYBPA2 did not complement the anthocyanin deficient phenotype of *A. thaliana pap1/pap2* mutant seedlings, but was able to activate *proAtANS* and *proAtDFR* when combined with *At*EGL3 in co-transfection experiments in *A. thaliana* protoplasts. Even though previous studies showed *AtEGL3* expression in *A. thaliana* seedlings (21), it is possible that the level was too low to activate anthocyanin biosynthesis in combination with *Musa*MYBPA2. Moreover, the At7 cell line used for co-transfection experiments was established more than 25 years ago. This long time of propagation in suspension cell culture caused a variety of genomic and transcriptomic changes (38). These changes could also explain differences between the analysis in seedlings and cell culture.

The expression profiles of *MusaMYBA1, MusaMYBA2* and *MusaMYBPA2* have previously been analysed in embryogenic cell suspension, seedling, root and different developmental stages of leaf, pulp and peel (30). The expression of *MusaMYBA1* was strongest in pulp (developmental stage S2-S3), *MusaMYBA2* expression was relatively low in all samples and *MusaMYBPA2* showed highest expression in seedlings and early developmental stages of pulp (S1). This gene expression pattern could give a hint at organ specific *MusaMYBA1, MusaMYBA2* and *MusaMYBPA2* activity in banana. Since flavonoid biosynthesis is largely regulated at the transcriptional level, it would be particularly interesting to analyse the expression levels of *MusaMYBA1, MusaMYBA2*, and *MusaMYBPA2* in anthocyanin rich organs such as bract or pseudostem to relate anthocyanin content to *MusaMYB* transcript abundance.

Deng *et al*. (32) performed an expression analysis of flavonoid biosynthesis related genes using leaves of banana plants overexpressing the anthocyanin repressor *MusaMYB4*. They found that the expression of *MusaMYBA1* and *MusaMYBPA2* is decreased along with *MusaDFR* and *MusaANS* in plants overexpressing *Musa*MYB4, compared to wildtype. This data supports a proposed functionality of *Musa*MYBA1 and *Musa*MYBPA2 as transcriptional activators of anthocyanin biosynthesis in banana, as *MusaMYB4* could cause a feedback regulation of the positive regulators of anthocyanin biosynthesis, of *MusaMYBA1* and *MusaMYBPA2*.

Very recently, Rajput *et al*. (33) showed that *Musa*MYBPA2 can activate the banana *ANS, ANR* and LAR promoters. Moreover, they showed that *Musa*MYBPA2 can partially rescue the proanthocyanin deficiency of the *A. thaliana tt2-1* mutant. The reported ability of *Musa*MYBPA2 to activate *proMusaANS* supports the proposed role of *Musa*MYBPA2 in the regulation of anthocyanin biosynthesis. However, the partial complementation of the proanthocyanin deficient phenotype of the *A. thaliana tt2-1* R2R3-MYB mutant, as well as the ability to activate the banana *ANR* and *LAR* promoters, shows an additional role in the regulation of proanthocyanidin biosynthesis. Such dual roles in the regulation of flavonoid biosynthesis have previously been suggested for R2R3-MYB transcription factors from grape, blueberry and apple (39-42). In several species, including *A. thaliana, Z. mays* and *Petunia hybrida*, anthocyanin biosynthesis is regulated by an MBW complex (21, 28, 43-45). Our co-transfection assays showed that *Musa*MYBA1, *Musa*MYBA2 and *Musa*MYBPA2 are able to activate *proAtANS* and *proAtDFR* as part of an MBW complex with *Zm*R and *At*TTG1 *in planta*. As presented in a previous study by Pucker *et al*. (30), it is known that *Musa*MYBA1, *Musa*MYBA2, and *Musa*MYBPA2 all contain a bHLH-binding consensus motif (27). Thus, the regulation of anthocyanin biosynthesis by *Musa*MYBA1, *Musa*MYBA2, and *Musa*MYBPA2 likely depends on bHLH and WD40 proteins. Based on our tree (Figure 2B), the bHLH encoding genes Ma05_g23010, Ma11_g19640, Ma11_g16060, Ma08_g11190 and Ma11_g03740 could be suitable candidates for further studies on the transcriptional activation of anthocyanin biosynthesis by MBW complexes. Moreover, future analyses should additionally include *Musa*TTG1 to identify a whole functional MBW complex in banana and to find out if the use of endogenous WD40 protein further improves the activation potential of the MBW complex. The presented results show that *Musa*MYBA1, *Musa*MYBA2, and *Musa*MYBPA2 have the capacity to transcriptionally activate structural anthocyanin biosynthesis genes in an MBW complex with a suitable bHLH partner. This is a step towards decoding the MBW complex-mediated transcriptional activation of flavonoid biosynthesis in banana. Moreover, it is a basis for further research towards increased anthocyanin production in banana which could promote fruit quality and disease resistance.

## Limitations

The co-transfection assays revealed that *Musa*MYBA1, *Musa*MYBA2, and *Musa*MYBPA2 can activate two structural genes of anthocyanin biosynthesis from *A. thaliana* as part of an MBW complex. In the future this analysis could be expanded to further genes. Moreover, co-transfection experiments with the promoters of structural anthocyanin biosynthesis genes and bHLH and WDR candidates from banana should be performed. In order to elucidate the regulatory role in banana and confirm possible target genes, future studies should include an overexpression of the three *Musa*MYBs in banana.

## Declarations

### Authors’ Contributions

MB, BP and RS planned the experiments. MB performed the experiments and analysed the data. RS and BW supervised the project. MB wrote the initial draft. RS and BP revised the manuscript.

## Acknowledgements

We are grateful to Melanie Kuhlmann and Janina Grabowski for their excellent technical assistance and to Andrea Voigt for her competent help in the greenhouse. We thank Martin Hülskamp for providing the *Zm*R entry clone plasmid and Ashutosh Pandey for providing the banana cDNAs.

## Abbreviations

ANS: anthocyanidin synthase
*A. thaliana*: *Arabidopsis thaliana*
bHLH: basic helix-loop-helix
C1: COLOURED ALEURONE1
DFR: dihydroflavonol 4-reductase
EGL3: ENHANCER OF GLABRA 3
Foc: *Fusarium oxysporum* f. sp. *cubense*
*M. acuminata*: *Musa acuminata*
*O. sativa*: *Oryza sativa*
PAP: production of anthocyanin pigment
TR4: tropical race 4
TTG1: TRANSPARENT TESTA GLABRA 1
*Z. mays*: *Zea mays*

## Supplementary information

Figure S1: Expression analysis of *MusaMYBA1, MusaMYBA2 and MusaMYBPA2* in *A. thaliana pap1/pap2* seedlings.

Figure S2: Full-length gels from Figure S1.

Table S1: Oligonucleotide primers used in this work.

Table S2: IDs of protein sequences used for the construction of the phylogenetic tree.

File S2: Methods

