## Supplementary information for "Three R2R3-MYB transcription factors from banana (*Musa* spp.) activate structural anthocyanin biosynthesis genes as part of an MBW complex"

**Supplementary material**

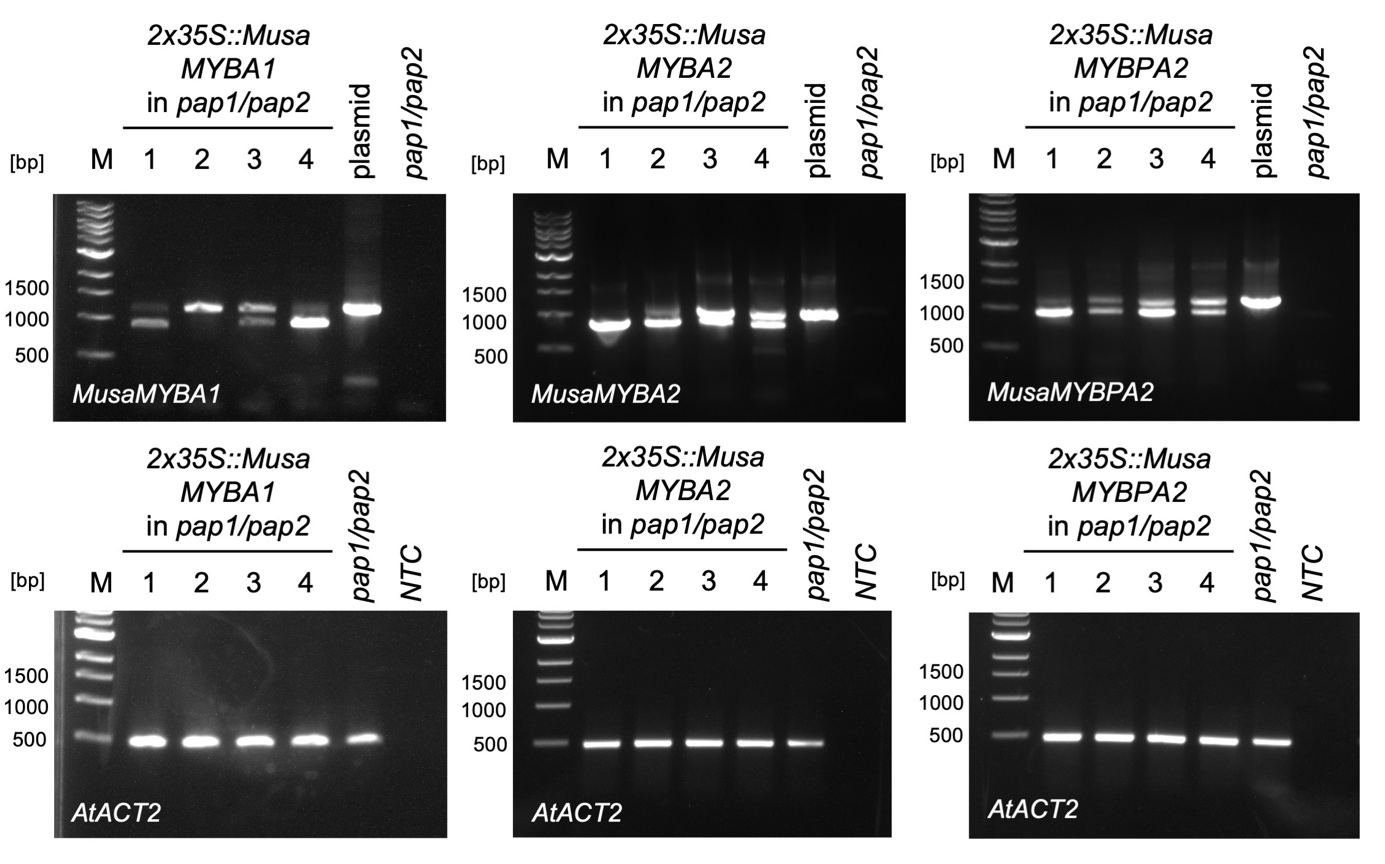

**Figure S1: Expression analysis of *MusaMYBA1, MusaMYBA2 and MusaMYBPA2* in *A. thaliana pap1/pap2* seedlings**. RNA was isolated from seedlings using NucleoSpin RNA Plant (Macherey-Nagel) according to the manufacturers’ instructions without DNase digestion to obtain both, RNA and genomic DNA. cDNA synthesis was performed from 1 μg nucleic acids using the ProtoScript First Strand cDNA Synthesis Kit (New England Biolabs). PCRs were run using a homemade Taq polymerase and standard protocols. Oligonucleotide primers are listed in Table S1. 1 kb DNA Ladder (New England Biolabs) was used as a standard (M). Different numbers represent individual transgenic lines and correspond to the lines shown in Figure 2. The binary plasmid pLEELA contains an intron which is spliced in *A. thaliana*. Thus, the two different bands in the *MusaMYB* specific PCRs correspond to genomic DNA (upper band) and cDNA (lower band).

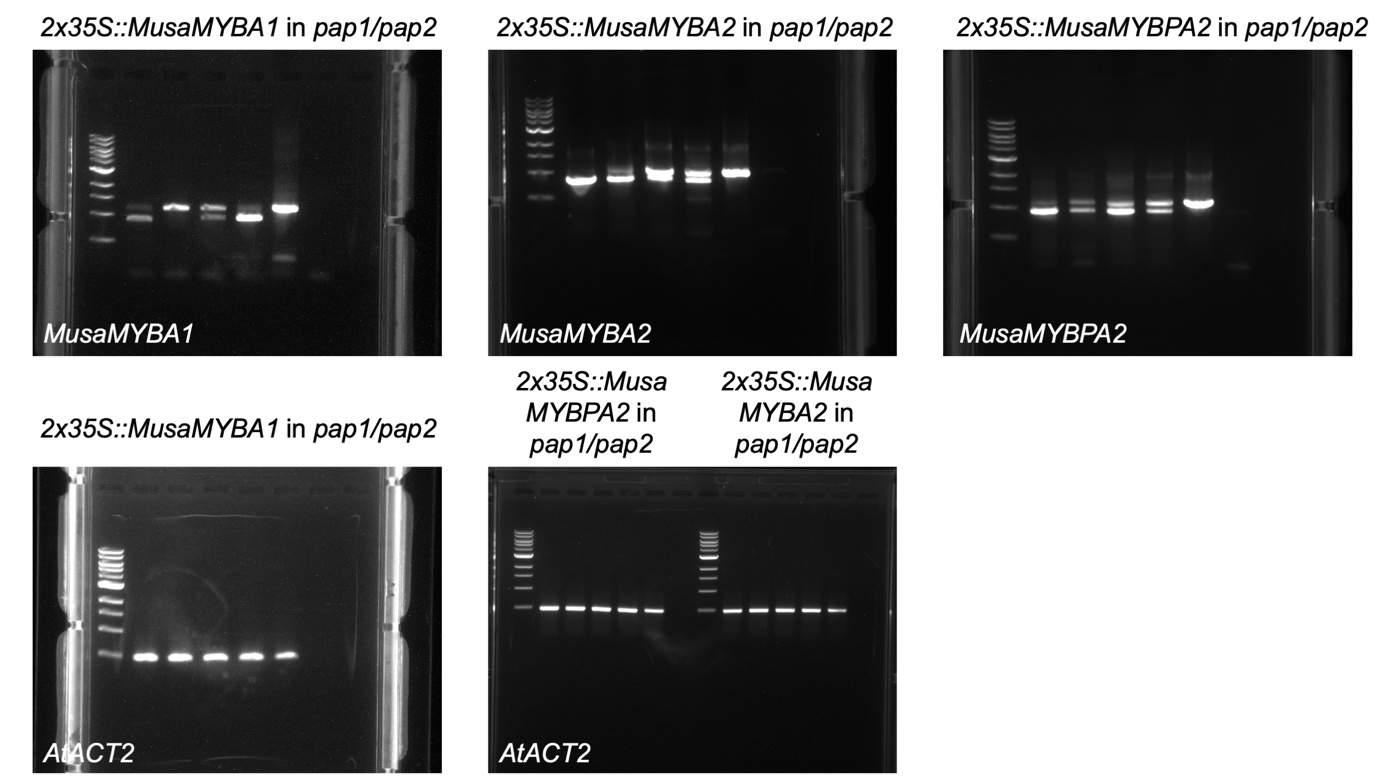

**Figure S2: Full-length gels from Figure S1.**

**Table S1: Oligonucleotide primers used in this work.**

| **Name** | **Sequence (5’-3’)** | **Description** |
| --- | --- | --- |
| RSt469 | TCCGCTCTTTCTTTCCAAGCTCAT | fwd Primer for *AtACT2* |
| RSt470 | TCCAGCACAATACCGGTTGTACG | rev Primer for *AtACT2* |
| RSt1516 | CCGTTCCATGGGCTAGAAGCTTCTCCTCC | fwd Primer in first exon of pLEELA |
| RSt1565 | attB1-ccATGGGAAGGAAGCCATGCTGTGCAAAGG | fwd Primer with attB1 for *MusaMYBA1* (Ma06_ 05960) CDS |
| RSt1566 | *attB2*-AACAACTTGGTGTCCAGTTCAGCCAGTCG | rev Primer with attB2 for *MusaMYBA1* (Ma06_ 05960) CDS |
| RSt1579 | *attB1*-ccATGGGAAGGAAGCCTTGTTGTTCAAAGG | fwd Primer with attB1 for *MusaMYBPA2* (Ma10_17650) CDS |
| RSt1580 | *attB2-*AGCTCCAATACCCCTGTTCGTCAGCTTC | rev Primer with attB2 for *MusaMYBPA2* (Ma10_17650) CDS |
| RSt1581 | *attB1-*ccATGGGAAGGAAGCCATGCTGCGTAAGGG | fwd Primer with attB1 for *MusaMYBA2* (Ma09_27990) CDS |
| RSt1582 | *attB2*-AGGTCCAATTCAGCCACTCCTCCGTATC | rev Primer with attB2 for *MusaMYBA2* (Ma09_27990) CDS |

attB1: GGGGACAAGTTTGTACAAAAAAGCAGGCT, attB2: GGGGACCACTTTGTACAAGAAAGCTGGGT

**Table S2:** **IDs of protein sequences used for the construction of the phylogenetic tree.**

| **MYB** |  |  |  | **bHLH** |  |  |
| --- | --- | --- | --- | --- | --- | --- |
| species | name | identifier |  | species | name | identifier |
| *A. thaliana* | PAP1 | AAG42001 |  | *Ma11_19640* |  | Ma11_19640 |
| *A. thaliana* | MYB114 | NP176812. 1 |  | *Ma05_23010* |  | Ma05_23010 |
| *A. thaliana* | PAP2 | NP176813. 1 |  | *Ma11_16060* |  | Ma11_16060 |
| *A. thaliana* | MYB113 | NP176811. 1 |  | *Ma11_3740* |  | Ma11_3740 |
| *A. majus* | VENOSA | ABB83828 |  | *Ma08_11190* |  | Ma08_11190 |
| *A. majus* | ROSEA1 | ABB83826 |  | *Ma06_24180* |  | Ma06_24180 |
| *V. vinifera* | MYBA1 | BAD18977 |  | *Ma03_18660* |  | Ma03_18660 |
| *P. hybrida* | AN2 | AAF66727 |  | *O. sativa* | Rb | MK636606 |
| *S. lycopersicum* | ANT1 | NP1234417 |  | *N. sylvestris* | AN1 | HQ589210 |
| *M. domestica* | MYB10 | ACQ45201 |  | *G. hybrida* | GMYC1 | AJ007709 |
| *D. variabilis* | MYB1 | AB601003 |  | *N. tomentosiformis* | AN1 | HQ589211 |
| *G. hybrid* | MYB10 | CAD87010. 1 |  | *Z. mays* | Lc | P13526 |
| *A. cepa* | MYB1 | KX785130 |  | *F. ananassa* | bHLH3 | JQ989284 |
| *A. thaliana* | MYB12 | AF062864 |  | *L. japonicus* | TT8 | AB490778 |
| *A. thaliana* | MYB11 | NP191820. 1 |  | *N. tabacum* | JAF13a | KF305768 |
| *A. thaliana* | MYB111 | NP199744. 1 |  | *R. sativus* | TT8 | KY651179 |
| *B. vulgaris* | MYB12 | NP001289993. 1 |  | *I. purpurea* | IVS | AB252663 |
| *Ma05_23640* |  | Ma05_23640 |  | *N. tabacum* | JAF13b | KF298397 |
| *Ma08_10260* |  | Ma08_10260 |  | *Z. mays* | B | X57276 |
| *Ma02_00290* |  | Ma02_00290 |  | *L. japonicus* | GL3 | AB492284 |
| *Z. mays* | P | AAC49394 |  | *B. oleracea* | TT8 | GU219990 |
| *V. vinifera* | MYBF1 | FJ948477 |  | *Z. mays* | IN1 | AAB03841 |
| *A. cepa* | MYB29 | KX785133 |  | *P. frutescens* | MYC-RP | AB024050 |
| *O. sativa* | IF35 | BAB20661 |  | *M. domestica* | bHLH3 | NM1294049 |
| *Ma07_19880* |  | Ma07_19880 |  | *S. melongena* | bHLH117 |  |
| *Ma06_04210* |  | Ma06_04210 |  | *P. frutescens* | Myc-F3G1 | AB103172 |
| *Ma07_19890* |  | Ma07_19890 |  | *L. hybrid* | bHLH1 | AB222075 |
| *A. thaliana* | MYB123 | NP198405. 1 |  | *P. hybrida* | JAF13 | AF020545 |
| *Ma03_28720* |  | Ma03_28720 |  | *N. tabacum* | AN1b | HQ589209 |
| *Ma03_07850* |  | Ma03_07850 |  | *O. sativa* | Ra | U39860 |
| *Ma09_15440* |  | Ma09_15440 |  | *O. sativa* | B2 | MK636610 |
| *Ma08_23390* |  | Ma08_23390 |  | *V. vinifera* | MYCA1 | EF193002 |
| *Ma03_07840* |  | Ma03_07840 |  | *Z. mays* | R | NM1111869 |
| *L. japonicus* | TT2a | AB300033 |  | *A. thaliana* | MYC1/bHLH012 | NM116272 |
| *T. aestivum* | MYB10 | AB599722 |  | *O. sativa* | Rc | XM15790874 |
| *Z. mays* | C1 | AAA33482 |  | *M. domestica* | TTL1 | NM1294049 |
| *O. sativa* | C1 | BAD04024 |  | *V. vinifera* | MYC1 | U447172 |
| *Ma10_17650* | MusaMYBPA2 | Ma10_17650 |  | *A. majus* | DELILA | AAA32663 |
| *Ma09_27990* | MusaMYBA2 | Ma09_27990 |  | *A. thaliana* | TT8 | NM117050 |
| *Ma06_05960* | MusaMYBA1 | Ma06_05960 |  | *B. napus* | TT8 | NM1315974 |
| *O. hybrid* | MYB1 | ABS58501 |  | *B. vulgaris* | cuct | XP019102724.1 |
| *V. hybrid* | MYB6 | ADQ57817 |  | *S. tuberosum* | AN1 | JX848660 |
| *A. thaliana* | MYB56 | NP197282. 1 |  | *G. triflora* | bHLH1 | AB459661 |
| *H. vulgare* | ANT | AKU38977 |  | *P. hybrida* | AN1 | AF260919 |
| *H. vulgare* | C1 | AK359777 |  | *A. thaliana* | EGL3 | NM105042 |
| *H. vulgare* | PL1 | AK358937 |  | *S. lycopersicum* | AN1 | XM19215815 |
| *P. equestris* | MYB11 | AIS35928 |  | *A. thaliana* | GL3 | NM148067 |
| *P. schilleriana* | UMYB6 | FJ039860 |  | *I. nil* | IVS | AB232775 |
| *D. kaki* | MYB2 | BAI49719 |  | *N. tabcaum* | AN1a | HQ589208 |
| *D. kaki* | MYB4 | BAI49721 |  | *M. domestica* | bHLH33 | DQ266451 |
| *P. trichocarpa* | MYB134 | XP2308528 |  | *L. hybrid* | bHLH2 | AB222076 |
| *T. arvense* | MYB14 | AFJ53053 |  | *B. vulgaris* | reum | XP010684257.1 |
| *V. vinifera* | MYBPA2 | ACK56131 |  |  |  |  |
| *A. majus* | MYB308 | P81393. 1 |  |  |  |  |
| *A. thaliana* | MYB0 | NP189430. 1 |  |  |  |  |
| *A. thaliana* | MYB23 | NP198849. 1 |  |  |  |  |
| *A. thaliana* | MYB3 | NP564176. 2 |  |  |  |  |
| *A. thaliana* | MYB32 | NP195225. 1 |  |  |  |  |
| *A. thaliana* | MYB4 | NP195574. 1 |  |  |  |  |
| *A. thaliana* | MYB66 | NP196979 |  |  |  |  |
| *A. thaliana* | MYB7 | NP179263. 1 |  |  |  |  |
| *F. ananassa* | MYB11 | JQ989282 |  |  |  |  |
| *F. ananassa* | MYB9 | JQ989281 |  |  |  |  |
| *F. tataricum* | MYB1 | JF313345 |  |  |  |  |
| *F. tataricum* | MYB2 | JF313347 |  |  |  |  |
| *M. truncatula* | PAR | HQ337434 |  |  |  |  |
