## Supplementary File 2 for "Three R2R3-MYB transcription factors from banana (*Musa* spp.) activate structural anthocyanin biosynthesis genes as part of an MBW complex"

**Methods**

**Cladistic analysis**

The approximately maximum-likelihood trees of plant MYB domain sequences and bHLH protein sequences were constructed as described previously (1): Multiple protein sequence alignments were created using MAFFT v7 (2) with default settings and cleaned with pxclsq (3). The tree was constructed with FastTree v2.1.10 using the WAG+CAT model (4) and visualised with interactive tree of life (5). GenBank IDs of protein sequences used for the construction of the tree are given in Supplementary Table S2.

**Plant material**

Banana plants (*M. acuminata* (AAA Group) ‘Grand Naine’) for RNA extraction were grown in the field in Lucknow, India. *Musa* gene annotation identifiers refer to the study of Martin *et al.* (6). *A. thaliana* Columbia‑0 (Col‑0, NASC ID N1092) and Nössen‑0 (Nö‑0, NASC ID N3081) accessions were used as wildtype controls corresponding to the *pap1/pap2* (*pap1*: Nö-0 background, PST16228; *pap2*: Col-0 background, SALK093731; (7)) double mutant, used for complementation assays. Hypocotyl-derived, cultured *A. thaliana* At7 cells (8) for co-transfection experiments were grown in MS medium (9) and maintained at 26 °C in the dark on a rotary shaker (105 rpm). Cells were subcultured weekly and cultures for protoplast isolation were inoculated 5 days prior to harvest.

**cDNA synthesis and molecular cloning**

RNA from Grand Naine peel was isolated according to a protocol from Asif *et al*. (10). cDNA synthesis was performed from 1 μg total RNA using the ProtoScript First Strand cDNA Synthesis Kit (New England Biolabs, NEB). Full length coding sequences (CDSs) were amplified using Q5® High-Fidelity DNA polymerase (NEB) and gene-specific primers (Supplementary Table S1). To generate Gateway Entry plasmids, PCR products were recombined into pDONR™/Zeo (Invitrogen) with BP clonase (Invitrogen) according to the manufacturer's protocol. Full length CDSs were introduced into the binary 2x35S overexpression vector pLEELA (11) and the 2x35S overexpression vector pBTdest (12) using Gateway LR reaction (Invitrogen). All plasmids were verified by Sanger sequencing.

***Agrobacterium tumefaciens*-mediated transformation of *A. thaliana***

*Agrobacterium tumefaciens* (GV3101::pMP90RK (13)) mediated gene transfer was carried out using the floral dip method (14). T-DNA integration from pLEELA-based plasmids into the *A. thaliana* genome was screened by BASTA selection and PCR-based genotyping (refer to Supplementary Figure 1 and Supplementary Table 1 for details).

**Determination of anthocyanin content**

*A. thaliana* seedlings were grown with 16 h of light illumination per day on 0.5x MS agar plates with 4 % sucrose to induce anthocyanin production. Anthocyanin content in a bulk of six-day-old seedlings was quantified as described by Mehrtens *et al.* (15). All samples were measured in three independent biological replicates. Error bars indicate standard deviation of the average (mean) anthocyanin content.

**Co-transfection experiments in *A. thaliana* protoplasts**

Cell culture, protoplast isolation, PEG-mediated DNA transfer into protoplasts and determination of standardised GUS activity were performed as described by Hartmann *et al*. (16). As an adaption we used 1.3 % (w/v) Cellulase Onozuka RS (Duchefa) and 0.8 % (w/v) Macerozyme R-10 (Duchefa) in 240 mM CaCl_2_ as the enzyme solution for protoplast preparation and 30 µg of plasmid DNA for the PEG-mediated DNA transfer into protoplasts. This comprised 10 μg of the promoter-GUS reporter construct, 0.5 μg of each 2x35S:transcription factor effector construct, 5 μg of the standardisation plasmid (pBT8 UBI-LUCm3), and an inactive luciferase expression vector (pBTΔLUC) to fill up to the amount of 30 μg of plasmid DNA. GUS-fused promoters, 520 bp *proAtDFR* and 1.95 kb *proAtANS,* as well as expression constructs for *At*PAP1 (*At*MYB75), *At*EGL3 (*At*bHLH2), *At*TTG1 and *Zm*C1 were available from previous studies (7, 12, 17). A *Zm*R entry clone (18) was kindly provided by Martin Hülskamp. Statistical analysis was performed using the Mann-Whitney *U* test (19).
